# Minicells as a potential chassis for engineering lineage-agnostic organisms

**DOI:** 10.1101/2020.07.31.231670

**Authors:** Eric Wei, Anton Jackson-Smith, Drew Endy

**Affiliations:** Department of Bioengineering, Stanford University, 443 Via Ortega, Stanford, CA, 94305; Department of Electrical Engineering, Stanford University, 350 Serra Mall, Stanford, CA 94305

**Keywords:** Minicells, genome engineering, organism engineering, synthetic cells, synthetic biology, expression capacity

## Abstract

**Background:** Genomes encode for organisms and thus genome synthesis implies the possibility of organismal synthesis, including the synthesis of organisms without constraint to lineage. Current genome-scale engineering projects are focused on minimization, refactoring, or recoding within the context of existing natural lineages. Minicells arise naturally as anucleate cells that are devoid of heritable genetic material but are capable of gene expression. Thus, minicells may serve as a useful starting point for developing lineage-agnostic organisms encoded by newly-designed synthetic genomes. However, the composition and expression capacity of minicells is fixed at the time of their formation. The possibility of reestablishing cellular growth and division starting from minicells and entirely heterologous synthetic genomes is unknown.

**Results:** We observed expression and segregation of functional proteins among mixed populations of reproducing cells and so-derived anucleate minicells via fluorescence microscopy. By adapting and integrating established methods of preparation and purification we were able to isolate minicells from a growing population of progenitor cells with a purity of at least 500 minicells per progenitor. We then used heterologous expression of plasmid-encoded green fluorescent protein to estimate the absolute expression capacity of minicells. We found that minicells can support the formation of 4.9 ± 4.6 × 10^8^ peptide bonds prior to exhausting their initial intrinsic expression capacity.

**Conclusions:** Minicells can be produced in large numbers with high purity and can also harbor and express engineered plasmids. The observed variation in minicell gene expression capacity suggests that about 13% of gene-expressing minicells can support the formation of more than one-billion peptide bonds, an amount sufficient to replicate known prokaryotic proteomes. Stated differently, while most minicells would require a sophisticated genetic ‘boot’ program to first increase minicell-specific expression capacity sufficient to instantiate newly reproducing lineages, a subset of minicells may be able to directly support whole genome, lineage-agnostic organism engineering.

## Background

Our increasing proficiency at engineering living matter is impacting a diversity of human activities including health, energy, agriculture, manufacturing, and our relationships with the environment [1–7]. One imperfect but operationally-useful measure of these impacts is the growth of the so-called bioeconomy, which has increased annually by ~10% in the United States over the past decade [8] and engages ~8% of the workforce in the European Union [9]. The increasing technological and economic impacts of bioengineering are undergirded by an increasing mastery of DNA design, from encoding single genes to complex pathways and cellular systems. More recently, capacities for designing, constructing, and handling DNA on the genomic scale [10,11] coupled with the dropping price and increasing fidelity of DNA synthesis and sequencing technologies [12] has made the building of functional synthetic genomes both technically practical and fiscally feasible. This recently emergent capacity to design and build genomes enables the pursuit of constructing, understanding, and operationally mastering the core unit of life – living cells.

Various approaches for building synthetic cells have made progress towards constructing functional synthetic genomes while also revealing gaps in our collective knowledge of how natural genomes encode viable cells. For example, top down genome minimization efforts (Fig. 1) have been used to determine one or more minimal gene sets capable of sustaining life [13–17]. As one specific example, work towards obtaining a minimal *Mycoplasma mycoides* genome generated the design of a 531 kbp genome (JCVI-syn3.0) encoding a functional set of 473 genes each individually essential for life yet for which 149 genes could not be assigned any specific biological function [13]. A more recent computational model of a JCVI-syn3.0-derived cell using flux balance analysis to categorize and quantify central-, nucleotide-, lipid-, cofactor-, and amino acid-metabolism was still unable to assign functions to 91 genes, at least 30 of which are essential [18]. The ongoing lack of a complete first-order understanding of all the genes essential for life presents a significant opportunity for the science of biology [19], while hindering the ability of engineers to operate at the cellular scale except via high-throughput tinker-and-testing.

**Figure 1.**
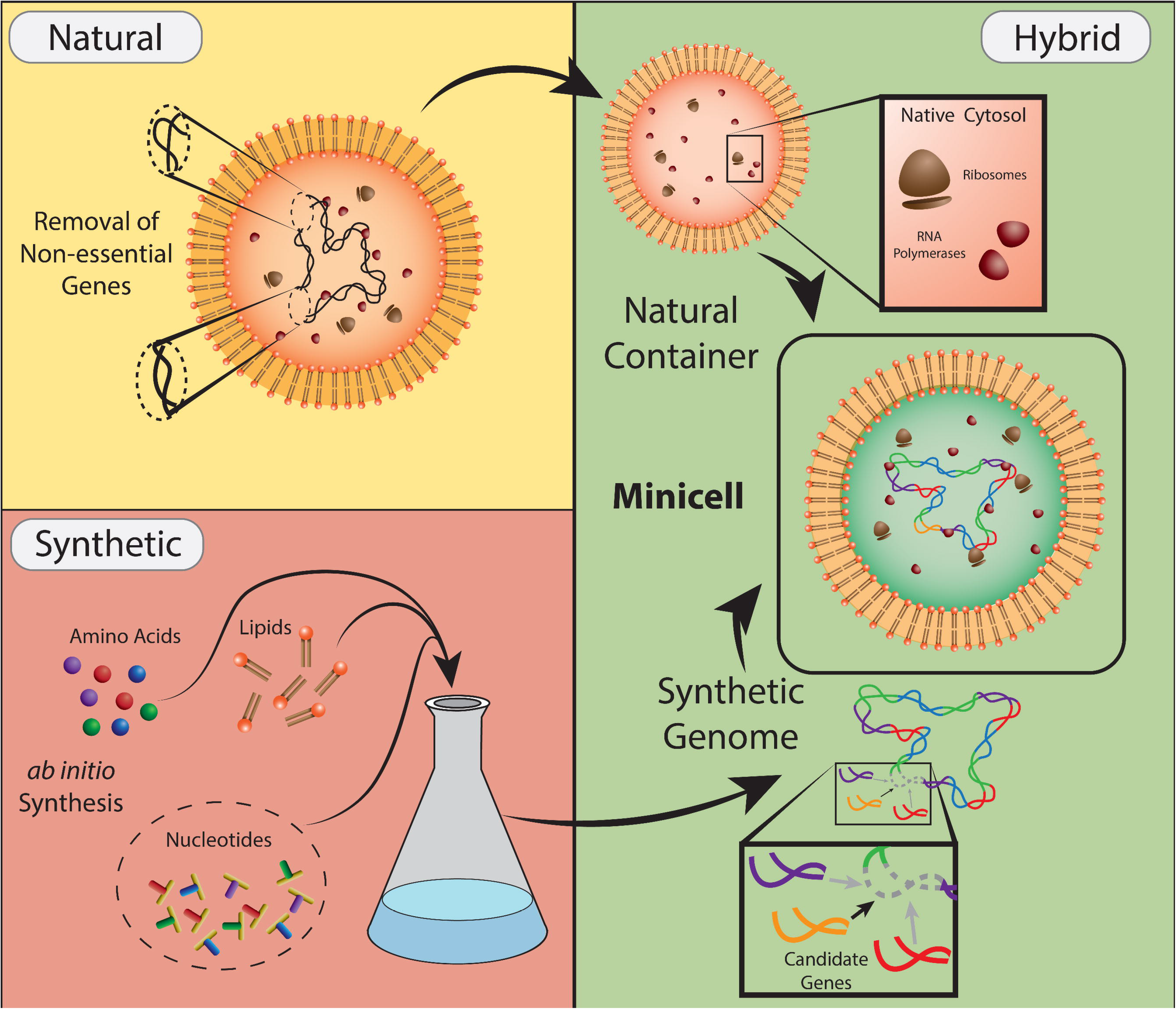
Minicells are a chassis housing existing natural expression machinery that could be used to express synthetic genomes and instantiate synthetic cells. Approaches for producing synthetic cells have traditionally applied top down minimization of genomes from removal of non-essential genes and bottom up synthesis of protocells from self-assembly macromolecules. The unanswered questions for building synthetic cells consist of identifying the unknown functions that are essential to life in the fully synthetic approach, and unknown genes of essential functions in the fully natural approach. Minicells combine the strengths of each by using the operational expression machinery and the container of natural cells and introducing fully synthetic genomes that are designed and engineered.

The dominant, complementary method for building synthetic cells utilizes bottom up additive approaches, combining from scratch the chemical and physical functions known or thought to be essential for life, with the aim of creating or recreating cellular life starting from lifeless ensembles of molecules (Fig. 1). Efforts along this track are focused on the co-assembly of artificial lipid membranes with cell extract-based or purified gene expression systems [21–24]. Additional work focused on the formation of minimal protocells as a model for abiogenesis strives to demonstrate the synchronization of genomic replication, often using RNA-based genomes and ribozymes, in unison with compartment division of droplets or unilamellar vesicles [25,26]. While significant progress has been made towards self-assembly, encapsulation, and synthesis and division of lipid membranes, the bottom-up creation of a full reproducing cell has not yet been realized. Moreover, it is unclear what may remain missing at the end of ongoing efforts; just as genome minimization efforts reveal essential genes of unknown function, we might also anticipate that bottom-up synthetic cell construction efforts will eventually reveal the complementary puzzle of essential functions that are unknown.

Given that the two dominant strategies for building fully-understood synthetic cells have not yet succeeded we sought to explore a third approach in which the strength of each existing method might complement the weakness of the other. One possibility would be to develop wholly-designed, lineage-agnostic genomes that, at least initially, operate within the molecular milieu derived from natural cells that have been depleted of all natural genetic material (Fig. 1). While the existence of containers for gene expression in the absence of expressible nucleic acids are vanishingly rare in nature, an example occurs in the form of minicells [27,28]. Minicells result from aberrant cell division events and contain a subset of proteins, RNA, and lipid membrane derived from a full cell while lacking genomic DNA. Bacterial minicell producing strains can occur from several different means [29,30], one example being an *E. coli* strain possessing a *minB* mutation. The absence of *minB* causes septation to appear at the poles during cell division resulting in spherical minicells ranging from 50 nm to a few microns in diameter (Figure 2D). Despite lacking a genome, minicells have been reported to possess the capacity for gene expression [31–35]. Although minicells themselves are lineage derived from their source organism, the possibility to express and replicate using an arbitrary genome that overtakes its endogenous componentry makes minicells a potentially attractive chassis for engineering synthetic cells. Stated differently and to summarize, top down and bottom up approaches help make clear our lack of understanding of essential genes of unknown functions and essential functions that are unknown; a “middle out” strategy starting from minicells might help bridge the gap.

**Figure 2.**
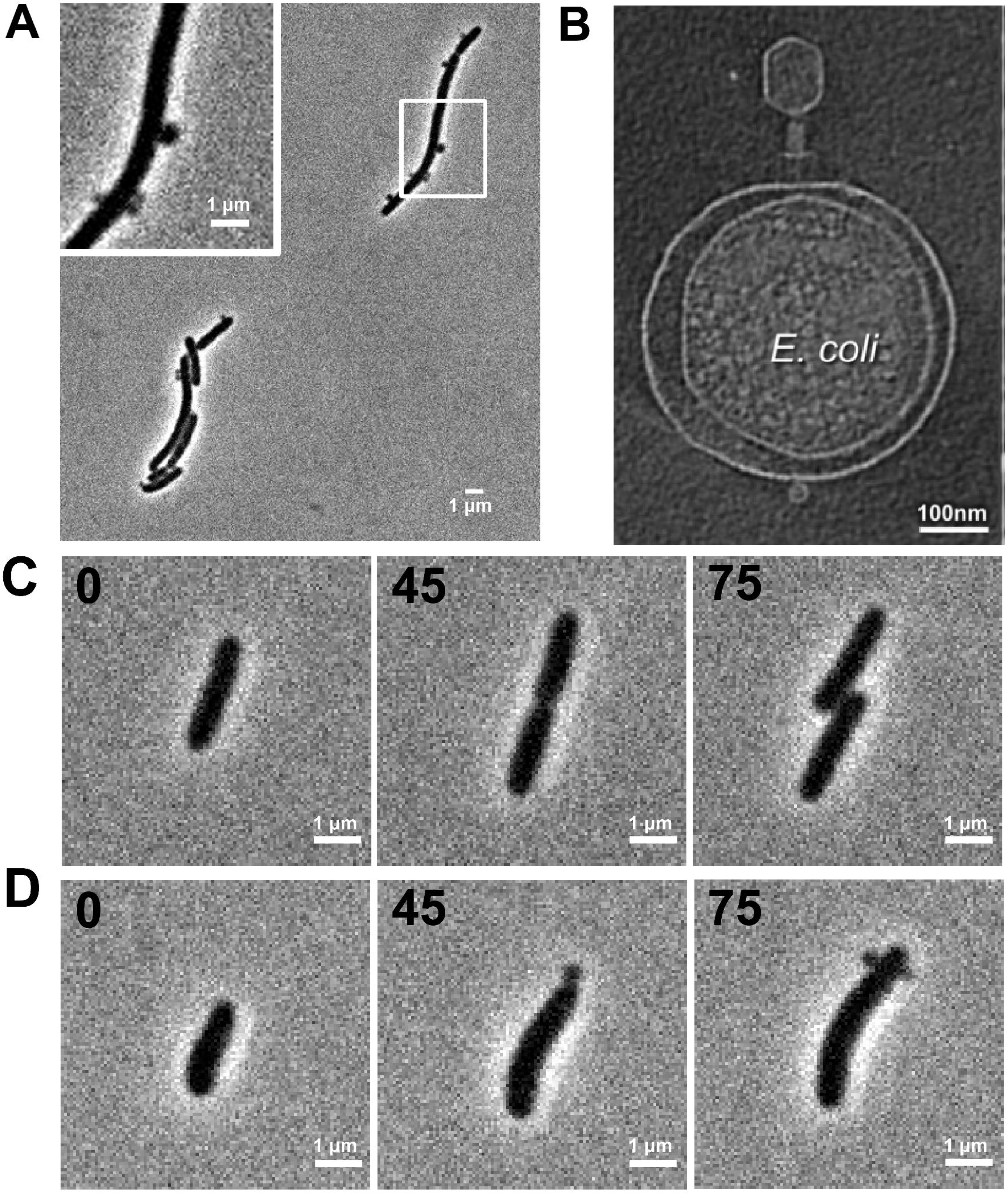
Minicells can be derived from mutant *E. coli* strains. (A) Phase contrast microscopy image of *E. coli* minicell producing strain DS410. (B) Electron cryotomography (Cryo-ET) slice of *E. coli* minicell infected with phage T4 (adapted from [39]). (C) Time lapse of single MG1655 (wildtype) cell and (D) DS410 at an elapsed time period of 0, 45, and 75 minutes.

Minicells have also found use in applications ranging from biosensing to drug delivery [35–39]. To expand minicell usage for prototyping synthetic cells, three technical questions will have to be addressed. First, are we able to reliably transfer heterologous DNA into minicells? Second, are we able to purify minicells from an initial growing culture of progenitor cells? Third, do individual minicells have the expression capacity necessary to express a genome encoding an entire living cell? Here, we establish working protocols and quantify the expression capacity of minicells, such that organism designers might better develop synthetic genomes to restore growth and reproduction of minicells without constraint to lineage.

## Results

### Heterologous proteins are present in newly-formed minicells, but where are they made?

Due to both their limited size and mechanism of formation, minicells are deficient in chromosomal DNA but receive a subset of cytosol, proteins, and membrane from their progenitor cells. To confirm this expectation, we used brightfield and fluorescence microscopy to observe minicell production from cells transformed with a plasmid that constitutively expresses green fluorescent protein (sfGFP) at medium copy number (10-20 copies per cell) (Fig. 3). While the progenitor cells continued to grow and divide, producing both viable progeny as well as minicells, the minicells themselves did not grow or divide (Fig. 3, 320’ brightfield panel). Most progenitor cells and at least some of the minicells harbored significant fluorescence (Fig. 3, middle and bottom rows).

**Figure 3.**
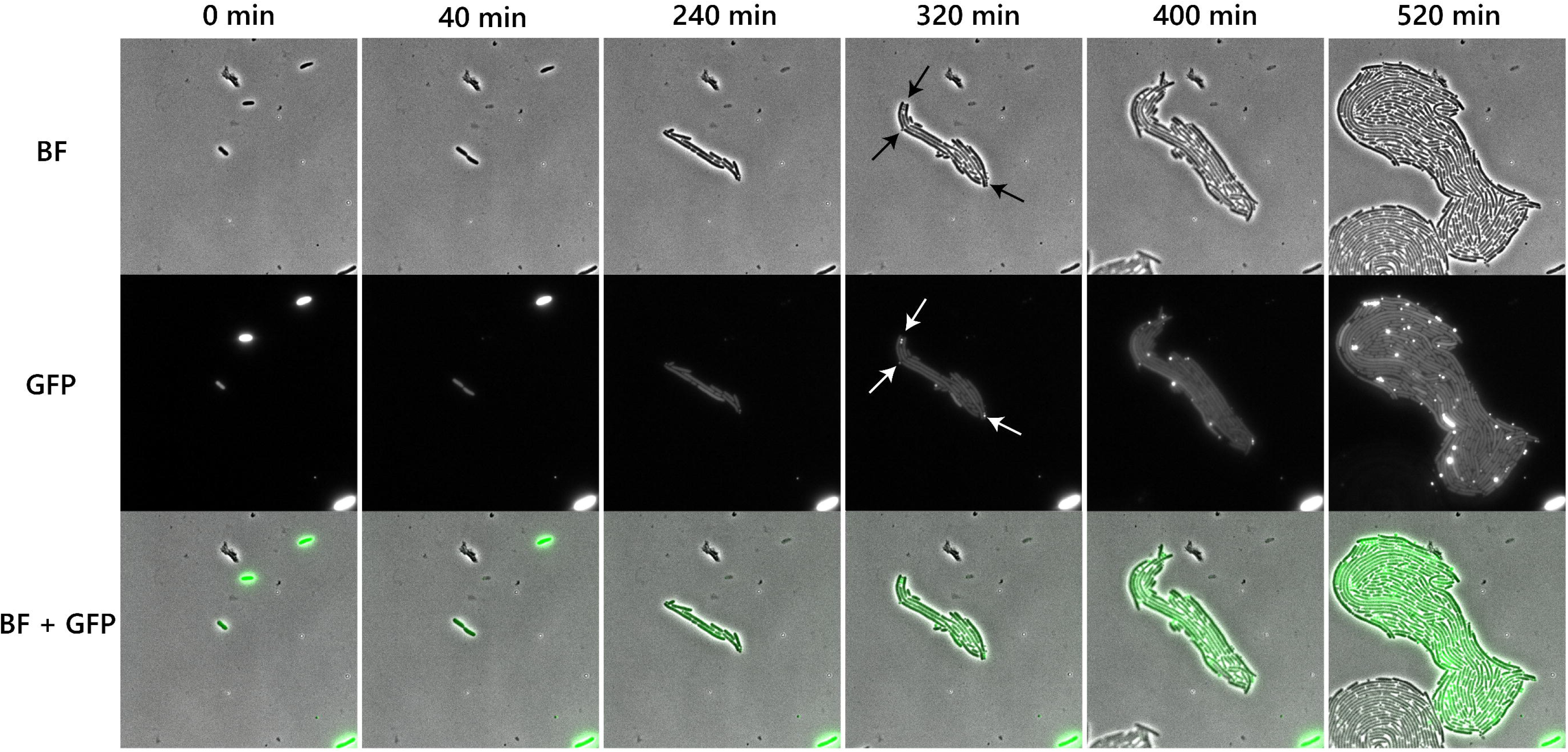
Newly-formed minicells harbor heterologous proteins from progenitor cells. **I**ndividual progenitor cells (strain DS410) bearing a constitutively expressed sfGFP plasmid for 520 minutes are captured using microscopy in the brightfield (BF) and GFP (490-510 nm) channels. Minicells (arrowed) are easily identifiable when examined under the GFP emission channel.

Minicell fluorescence typically intensified and appeared more highly concentrated than in progenitor cells. Upon formation, minicells receive all the components necessary for gene expression, and so those minicells that receive copies of the sfGFP-encoding plasmid have the possibility for continued expression of sfGFP. However, it is unclear whether the heightened fluorescence signal results from sfGFP production in the minicells themselves. The increased signal could be due to progenitor-cell expression of sfGFP followed by segregation of progenitor-expressed proteins into minicells that undergo post-translational maturation. To evaluate whether minicells are a suitable chassis for cell builders, we must better quantify to what extent minicells can express heterologous genes.

### Minicells can be extracted and purified from growing culture of progenitor cells

In order to better perturb and quantify minicells in the absence of progenitor cells, we sought to purify minicells with high yield and purity. As minicells themselves are unable to replicate, they are dependent on progenitor cells to produce an abundant number sufficient for characterization and downstream processing. Historically, differential separation of minicells has been conducted using sucrose gradients, filtration, and centrifugation methods that leverage the differing size and densities of minicells compared to their progenitors [27,40,41]. Alternatively, selection methods have been performed using penicillin-based antibiotics to inhibit synthesis of the peptidoglycan layer, exploiting the minicells’ arrested growth to lyse progenitor cells while leaving minicells intact [42]. We used an approach that combines differential centrifugation and antibiotic selection to purify minicells [43] and then quantified the purity and yield of the resulting preparations.

To start we grew cultures of the minicell-producing strain to late exponential growth phase (OD ~1), where a majority (~75%) of minicells within the mixture will have been formed within a window of two cell doublings. We then centrifuged down the mixed culture leaving most of the minicells within the supernatant (Fig. 4A). We observed via flow cytometry that two population clusters are separable when sorted by side scattering, signifying clusters of minicells at low scattering and progenitor cells at high scattering (Fig. 4B). Minicells represented 26.7% of cells that are pelleted, and a sizable number of progenitor cells are left in the supernatant and must still be removed (Fig. 4C). We then added ceftriaxone, a penicillin-based antibiotic, to lyse growing cells (Fig. 4D). By this purification process, we obtained a yield of 6 × 10^8^ minicells per 1 L of starting culture and a purity of 500 minicells per remaining viable progenitor cell, as measured by plate assay. The purity can be further increased through post-purification centrifugation of cell debris and FACS for minicell sorting and extraction (Fig. 4E).

**Figure 4.**
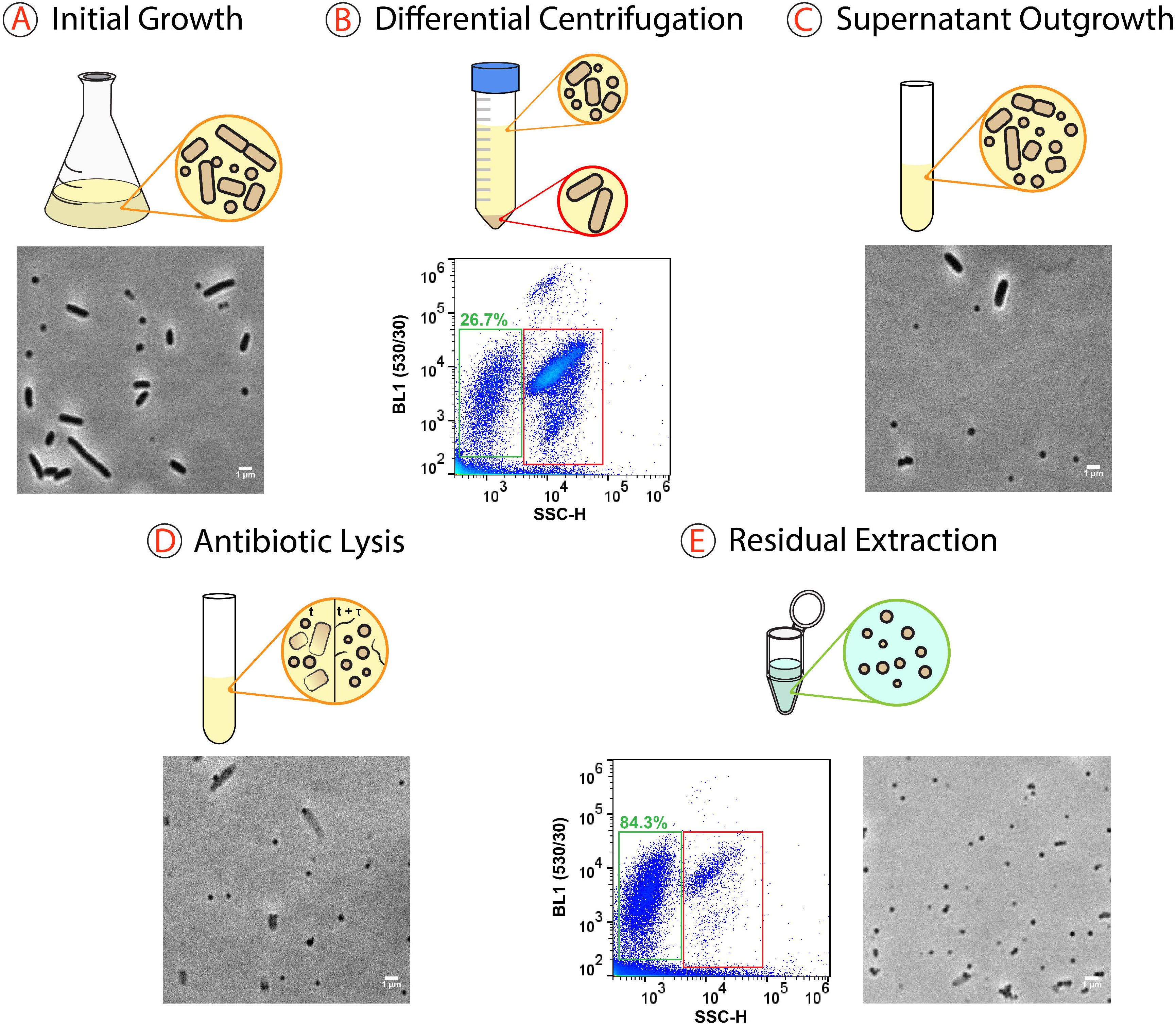
Purification yields large quantities of high-purity freshly-formed minicells. (A) An overnight culture of DS410 is backdiluted and grown to exponential growth phase (OD_600_ ~1). (B) We centrifuge down the culture pelleting most of the progenitor cells while leaving a population of minicells and small progenitor cells in the supernatant. Flow cytometry data reveals 26.7% of pellet contains potential minicells. (C) The supernatant is incubated to reinitiate growth of progenitor cells. (D) The addition of ceftriaxone causes the growing progenitor cells to lyse leaving minicells and cell debris. (E) Minicells can be extracted through methods such as FACS (sorting data pictured on left) or suspension in supernatant after centrifugation of cell debris (resultant minicells pictured on right).

To eliminate expression activity from the small number of remaining progenitor cells, we eluted the remaining minicells within sterile PBS and applied additional ceftriaxone. We conducted viability assays using dilution-based plating on growth media and saw no viable colonies when plating 2 × 10^9^ minicells incubated overnight at 37°C. We also saw no increase in optical density of the culture under sustained incubation at 37°C over two days. Using these methods, we can maintain quantities of minicells to utilize and characterize without any background presence of progenitor cells.

### Heterologous expression measured in bulk minicell preparations

To study whether the increased fluorescence observed over time is due to synthesis of new fluorescent proteins within minicells, we conducted an assay measuring translation activity in purified minicells. Previous evidence of gene expression within minicells have been reported in the forms of expression from heterologous plasmid DNA [31–35], producing transcription and translation activity from minicell lysate [41,44], or via susceptibility to phage infection and production of phage particles [45–49]. We sought to directly verify and quantify gene expression within minicells using a fluorescence-based assay.

We purified minicells as described from a minicell-producing *E. coli* strain harboring a sfGFP-expressing plasmid. We relied on plasmid segregation during cell division for introduction of heterologous DNA to minicells. A significant proportion of minicells should receive the plasmid as well as plasmid-bound RNA polymerase, thereby increasing minicell-based transcription [41,44]. We observed constitutive expression of sfGFP in minicells (Figure 5). We used a protein synthesis inhibitor (chloramphenicol) to distinguish between newly-expressed GFP and latent maturation of GFP made in progenitor cells. We measured an increase in fluorescence over time for minicell samples both with and without the translation inhibitor. However, after one hour, we observed no fluorescence accumulation in minicells exposed to the translation inhibitor, whereas untreated minicells accumulated roughly twice as much fluorescence, consistent with expression and maturation of new heterologous proteins inside minicells.

**Figure 5.**
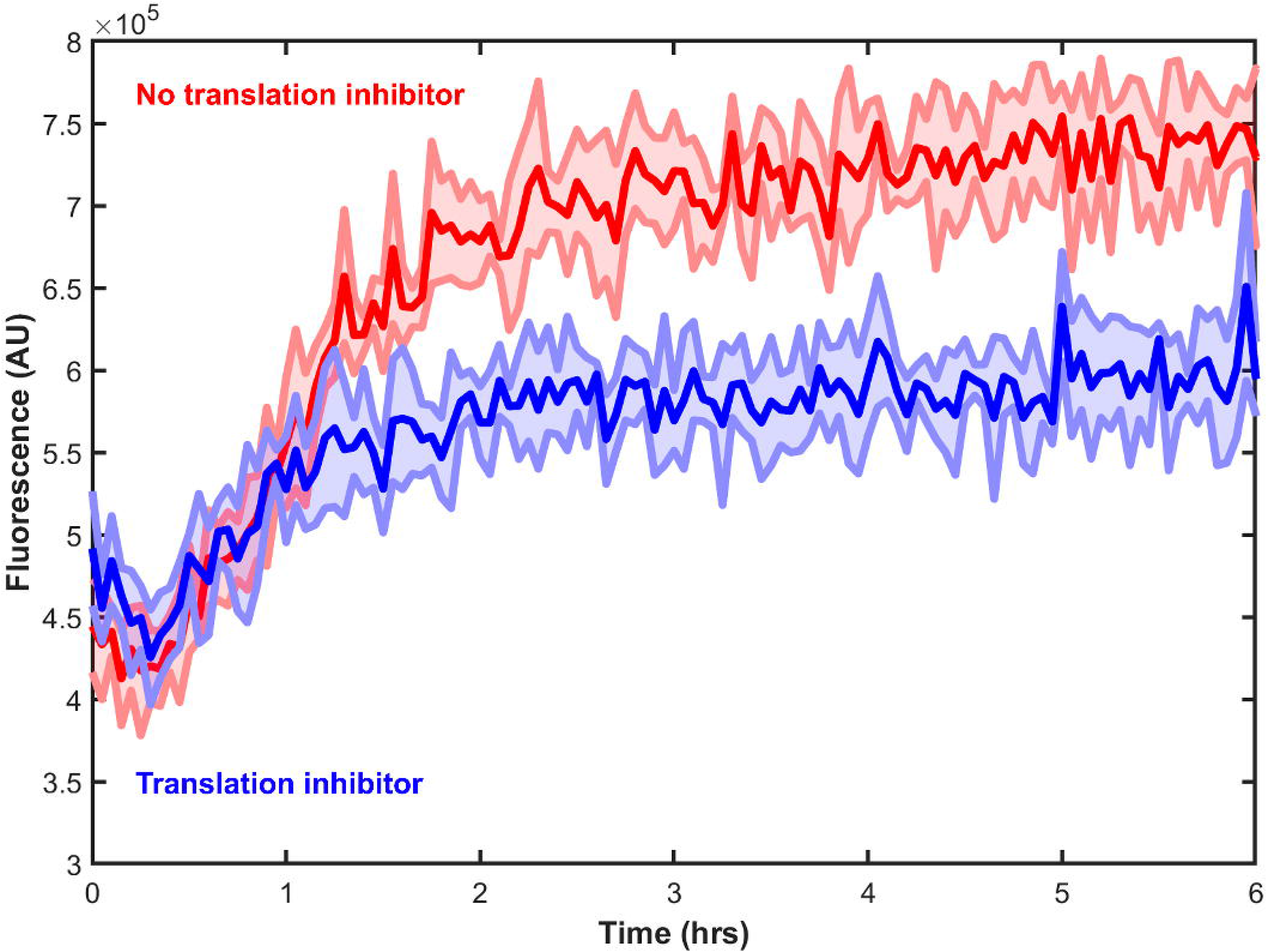
Heterologous gene expression occurs within minicells. Minicells are purified as previously described with and without the addition of translation inhibitor (chloramphenicol) applied post purification. Solid lines denote the mean and the shaded regions denote the standard deviation (n=3).

### A small fraction of individual minicells should be capable of expressing entire genomes

We next sought to understand the expression capacity of individual minicells. Due to the random nature of their formation, minicells are expected to possess high diversity in their composition and expression capacity. We first confirmed this notion by observing minicells via time-lapse fluorescence microscopy (Fig. 3), where the increase in fluorescence within individual minicells varied widely compared to neighboring progenitor cells and nearby minicells.

We then used time-lapse microscopy to track fluorescence production in individual minicells under two conditions: minicells formed immediately from their progenitor cells and minicells purified as described. We tracked individual progenitor cells and newly-formed minicells every 15 minutes (Fig. 6A). We wrote and used a script for tracking individual minicells throughout the movie and the fluorescence of minicells was recorded. Minicells recorded in this fashion have, on average, a nearly 3-fold increase in fluorescence over a two hour window. We tracked minicells prepared by our previously described purification process using the same methodology and filter settings (Fig. 6B) and found that the purified minicells have on average a 2.7-fold increase in fluorescence over time. We converted the measured fluorescence signal to an absolute number of sfGFP molecules produced (Supplementary Fig. 1) and estimate that purified minicells are able to form 4.9 ± 4.6 × 10^8^ peptide bonds. Compared to the ~1 × 10^9^ peptide bonds necessary for making an *E. coli* proteome at a doubling time of 30 minutes [50], at least 13% of gene-expressing minicells should be able to express a full microbial proteome.

**Figure 6.**
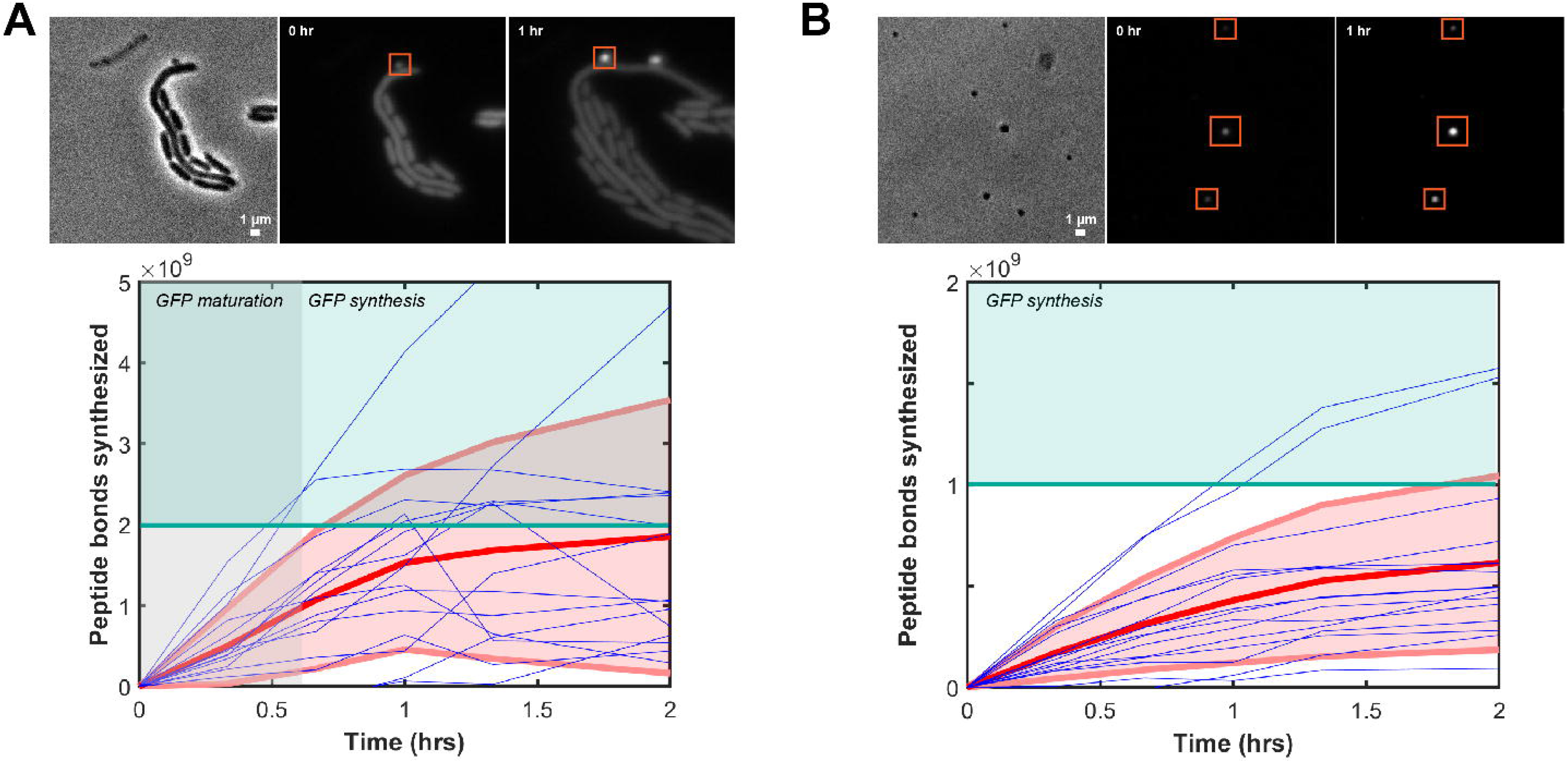
A small fraction of minicells appear to possess the expression capacity needed to instantiate synthetic cells. (A) Using time lapse videos of single cells of minicell producing strain DS410, minicells that arise from cell division events are tracked (time = 0) and fluorescence signal is tracked over time. 20 individual traces are presented in blue, and the mean and standard deviation of 100 traces are shown. The shaded region signifies the time required for 90% of sfGFP to complete maturation process *in vivo* [62]. The green regime denotes expression capacity necessary to express *E. coli* proteome at a doubling rate of 30 minutes. (B) Minicells containing sfGFP plasmid are tracked post-purification process (time = 0) from initial DS410 cells with sfGFP expressing plasmid. 20 individual traces are presented in blue, and the mean and standard deviation of 397 traces are shown. The green line denotes the expression capacity necessary to express the *E. coli* proteome.

## Discussion

Minicells possess several intrinsic qualities that make them an attractive option for prototyping genomes and constructing synthetic cells. Most notably, minicells provide an ensemble of functions necessary for life, including unknown functions, that may serve as a “crutch” for bottom-up cell building approaches; life-essential functions that are currently unknown may already exist within minicells. Stated differently, in contrast to liposomes and primitive protocells, minicells inherit known and unknown functional molecules that have already evolved to persist as a physically-structured ensemble. Thus, while minicells may be ill suited for understanding abiogenesis, their suite of evolved enzymes, polymerases, tRNAs, ribosomes, and other components form a complex milieu that could provide an advanced starting point for building and testing lineage-agnostic synthetic genomes. Additionally, although minicells themselves are unable to replicate, the progenitor cells that produce minicells can rbe eengineered in ways that vary minicell composition, as might be useful in studying or debugging genome-scale engineering projects.

Practically, working with minicells introduces other constraints that may limit their usefulness as chassis for synthetic cell building. For example, many traditional laboratory methods, especially those that rely on growth or selection of specific genotype or phenotype, cannot be used with non-growing minicells. As an example, we tested direct chemical transformation and electroporation of sfGFP-expressing plasmids into minicells but found no evidence of successful transformants (Supplementary Fig. 2). The lack of transformants could be explained by low efficiencies of transfer or by the lack of plasmid-bound RNA polymerase from the purified plasmid DNA used during transformation.

Another constraint of minicells is their reduced physical size and expression capacity that may, in turn, limit the size of genome that can initially be contained and expressed within most minicells. For example, using a volume of 1 nm^3^ per base pair of packed DNA, an entire 4.6 MB *E. coli* genome would occupy the entire volume of a 200 nm-diameter minicell. Smaller minicells might thus necessitate a “boot order” for the staged expression of core life functions prior to expressing all functions needed for growth and division. Addressing the challenge of booting up smaller minicells may be revealing for synthetic cell engineering, in general, as a goal in and of itself. However, we note that conjugation of genomic DNA into minicells followed by recovery of cell growth has been reported albeit as extraordinary low frequencies [51], suggesting that synthetic genomes may be able to be transferred to minicells via conjugation. Alternatively, the genome of progenitor cells could be degraded, leaving behind a maxicell (i.e., the functional equivalent of an anucleate minicell but with the size and resourcing of a normal-sized cell) [52–54].

Minicells also provide a means for cleanly instantiating *ab initio* genomes without constraint to specific natural lineages (Fig. 7). For example, if one is interested in minimal genomes it may be important to consider genome minimization beyond the context of a specific natural lineage. As a more specific example, the JCVI-syn3.0 genome uses the pre-existing *Mycoplasma mycoides* ATP synthase as its source of ATP. However, there exist multiple means for cells to produce or obtain ATP. If a potential synthetic cell was supplied with an ATP-rich environment then that cell might be able to more simply utilize an ATP/ADP translocase adapted from a bacterial endosymbiont [55] as its energy source – an impossibility if genome-scale researchers strictly adhere to lineage-specific genome minimization. The potential for construction by the means of composing and integrating functions across lineages could thus enable the process of building up of cells more generally.

**Figure 7.**
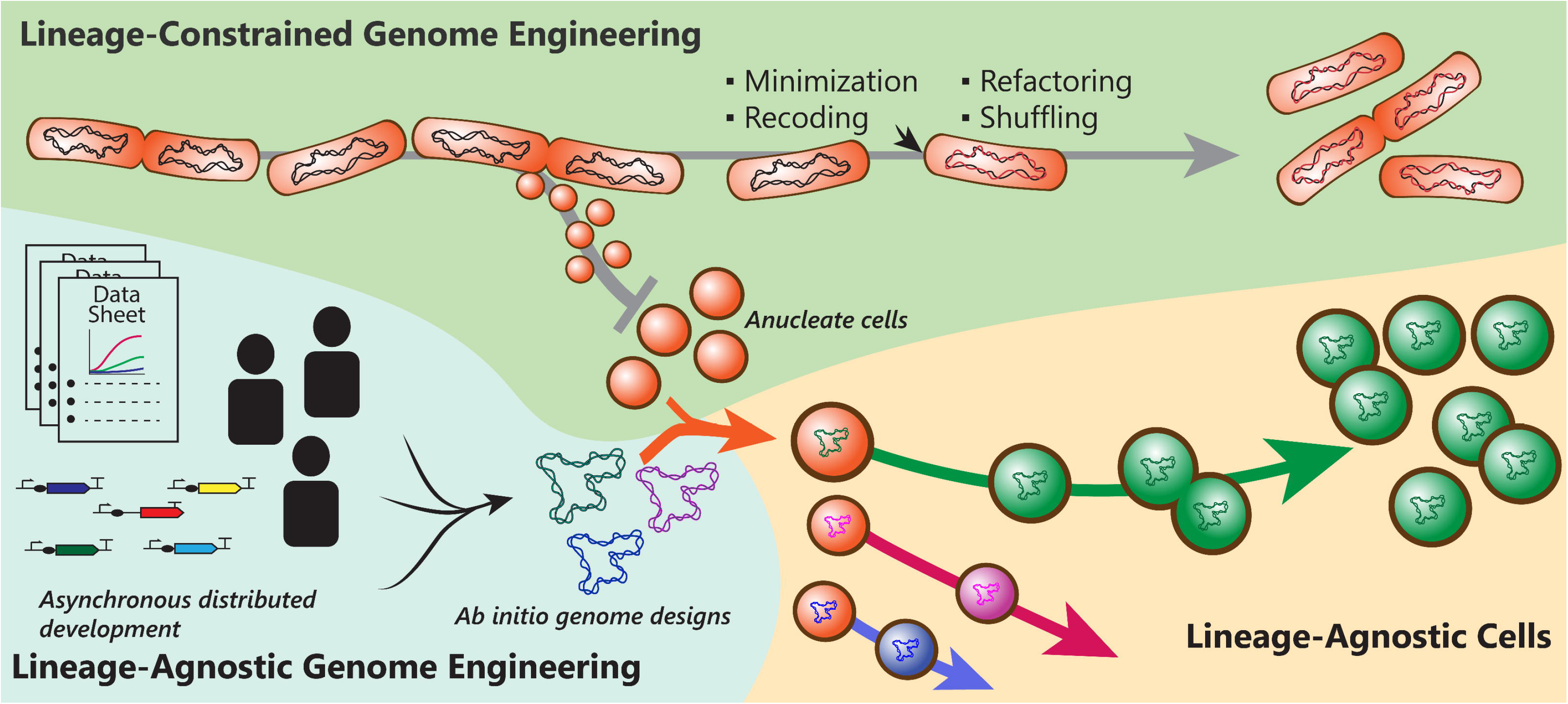
Enabling engineering of lineage-agnostic genomes could facilitate the distributed development and construction of operationally-understandable cells. Current methods for engineering cells and genomes involve iteration or re-assortment of genes from natural genomes (top) and are reliant upon an existing lineage for design and instantiation of new cell strains. The design of genomes *ab initio* (lower left) promote asynchronous distributed development without constraint from a single lineage. Successful implementation of designed genomes in membrane bound anucleate expression systems (lower right) permits the synthesis of novel, rationally-designed repurposed cells.

Building cells from scratch and without constraint to lineage will also require expertise across multiple areas of cell biology, biochemistry, biophysics, bioengineering, imaging, and many other disciplines. To best coordinate and integrate such work, we might take note of lessons from other engineering disciplines that have overcome systems-scale integration challenges. One example we take inspiration from is the development of the UNIX operating system [56]. In a similar vein to how the UNIX operating system was purposefully designed to enable communal development of software [56,57], we suggest that the emerging global network of cell builders (e.g., BaSyC, Build-A-Cell, et al.) might choose to design the process by which cells are built in ways that enable fellowship and collaboration; how we organize ourselves will impact how we organize our cells, and vice versa. Minicells or their equivalents, being an easily producible and distributable chassis with the means for executing *ab initio* lineage-agnostic genomes, have remarkable potential as a starting system for the coordinated work that will be needed to realize fully-understood synthetic cells.

Antoine de Saint Exupéry eloquently stated that “building a boat isn’t about weaving canvas, forging nails, or reading the sky. It’s about giving a shared taste for the sea, by the light of which you will see nothing contradictory but rather a community of love” [58]. Perhaps the greatest benefit the mission of rejuvenating minicells might provide is the vivid dream of the requisite knowledge, design, measurements, tools, and community that would be necessary to realize the goal of collaborative construction of synthetic cells.

## Conclusions

We developed methods for making and characterizing minicells for the purpose of enabling the engineering of lineage-agnostic organisms. We confirmed that minicells have the capacity to harbor and express user-designed genetic material. We showed with the use of translation inhibition and fluorescence microscopy that minicells are capable of ab initio protein synthesis and that minicell expression capacity can be quantified with single minicell resolution. Based on our observations a fraction of minicells so-produced should be able to express genes sufficient to instantiate a free-living microbe.

## Methods

### Strains and plasmids

Minicell producing strain DS410 and derivatives are obtained from Yale Coli Genetic Stock Center (CGSC). The sfGFP plasmid used to quantify expression in minicell strain was derived from previous work [59].

### Purification of minicells

We used a previous combination approach of antibiotic and size centrifugation [43] with some modification. Unless otherwise specified, we grew cultures at 37°C and 250 RPM. We cultured 5 mL culture overnights of DS410 in LB, with addition of 25 μg/mL chloramphenicol when using the sfGFP plasmid. We back-diluted the culture 1 mL into 1 L of fresh LB and grown for 6 hours, at the end of the exponential growth phase (OD ~1). We centrifuged the culture at 2000 × g for 10 minutes at 4°C and retained the supernatant. We centrifuged the supernatant at 10,000 × g for 10 minutes at 4°C. We resuspended the pellet in 50 mL of LB and incubated in shaker for 45 minutes. We added ceftriaxone to the culture at a final concentration of 100 μg/mL and continued incubating the culture for 2 additional hours. Afterwards, we centrifuged the culture at 400 × g for 5 minutes to removal of cell debris and the decanted supernatant is centrifuged at 10000 × g for 10 minutes. We resuspended the final pellet in 20 mL of PBS stored at 4°C.

### Quantification of yield and purity of minicell purification

We obtained minicell [43] and *E. coli* [60] cell counts using previously established formulas

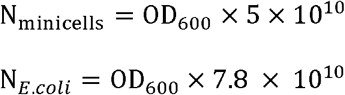

where N is the count per mL volume.

We assumed the portions of the cell culture post purification contributing to the OD_600_ reading consist solely of minicells, *E. coli* cells, and cell debris. We calculated the ratio of contribution of OD_600_ consisting of *E. coli* cells to cell debris using a non-minicell producing strain (MG1655), with the number of *E. coli* cells measured by plate assays and calculated the expected OD_600_ using the above formula. We subtracted this reading to the measured OD_600_ and attribute the rest of the OD_600_ to cell debris. This calculation on MG1655 calculated 1.12% of the OD_600_ of the purified minicell culture as *E. coli* cells. We ran the purified minicells through flow cytometry (Attune NxT) using fluorescence (490 nm excitation, 530 nm emission with 30 nm bandwidth) and SSC gating settings to establish counts within a minicell and large cell bin (Figure 4E). We categorized all counts within the gated region as *E. coli* and cell debris and compared the ratio to hits in the minicell bin (1:5). The concentration of minicells can be calculated by the OD_600_ of the purified minicells and carry-over progenitor cells through

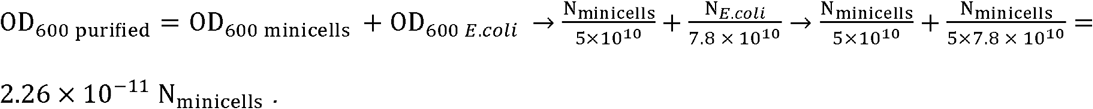

### Phase contrast and fluorescence microscopy

We used time lapse microscopy using an inverted light microscope (Nikon Eclipse TE-2000 E) with a fluorescent excitation lamp (Lambda XL, 470 nm excitation, 500 nm emission) and image capture software (MicroManager [61]). We seeded and stabilized cells and minicells using 1.5% low melting point agarose pads with LB media. We incubated cells and minicells at 37°C and acquired time lapse images every 40 minutes for 18-hours. The images and videos shown are identically leveled with a peak of 0.1% pixel saturation for the frame with the highest produced signal.

### Quantification of expression capacity of minicells

We grew and purified 1 L of minicell-producing culture with the sfGFP plasmid and 1 L of minicell-producing culture without the sfGFP plasmid and eluted into a final volume of 500 μL. We pipetted the minicell preparation into a 96 well plate (Greiner Bio-One M-8935) with 100 μL volume. We purified minicells without the sfGFP plasmid to subtract as background. For wells with translation inhibitor, we added chloramphenicol at a final concentration of 25 μg/mL. We used a plate reader (SpectraMax i3) to measure sfGFP fluorescence (485 nm excitation 10 nm bandwidth, 520 nm emission 10 nm bandwidth) every 5 minutes over an 8-hour time period at 37°C.

### Quantification of expression capacity of minicells

We established a standard curve using 6×His-purified sfGFP linking fluorescence measured on the plate reader and concentration of sfGFP (Nanodrop). We calculated the number of GFP molecules present in each well using the standard curve. We then equate the increase in fluorescence of bulk minicells in the plate reader (Figure 5) to the expression of number of sfGFP molecules. We analyzed images taken from the fluorescence microscope time lapse (Figure 3 and Figure 6) to attribute the proportion of fluorescence increase of the bulk mixture to that of sfGFP molecules expressed by individual minicells. We utilize the number of sfGFP molecules produced (240 amino acid length) to quantify the expression capacity as the number of peptide bonds formed.

---

**Supplementary Figure 1.**
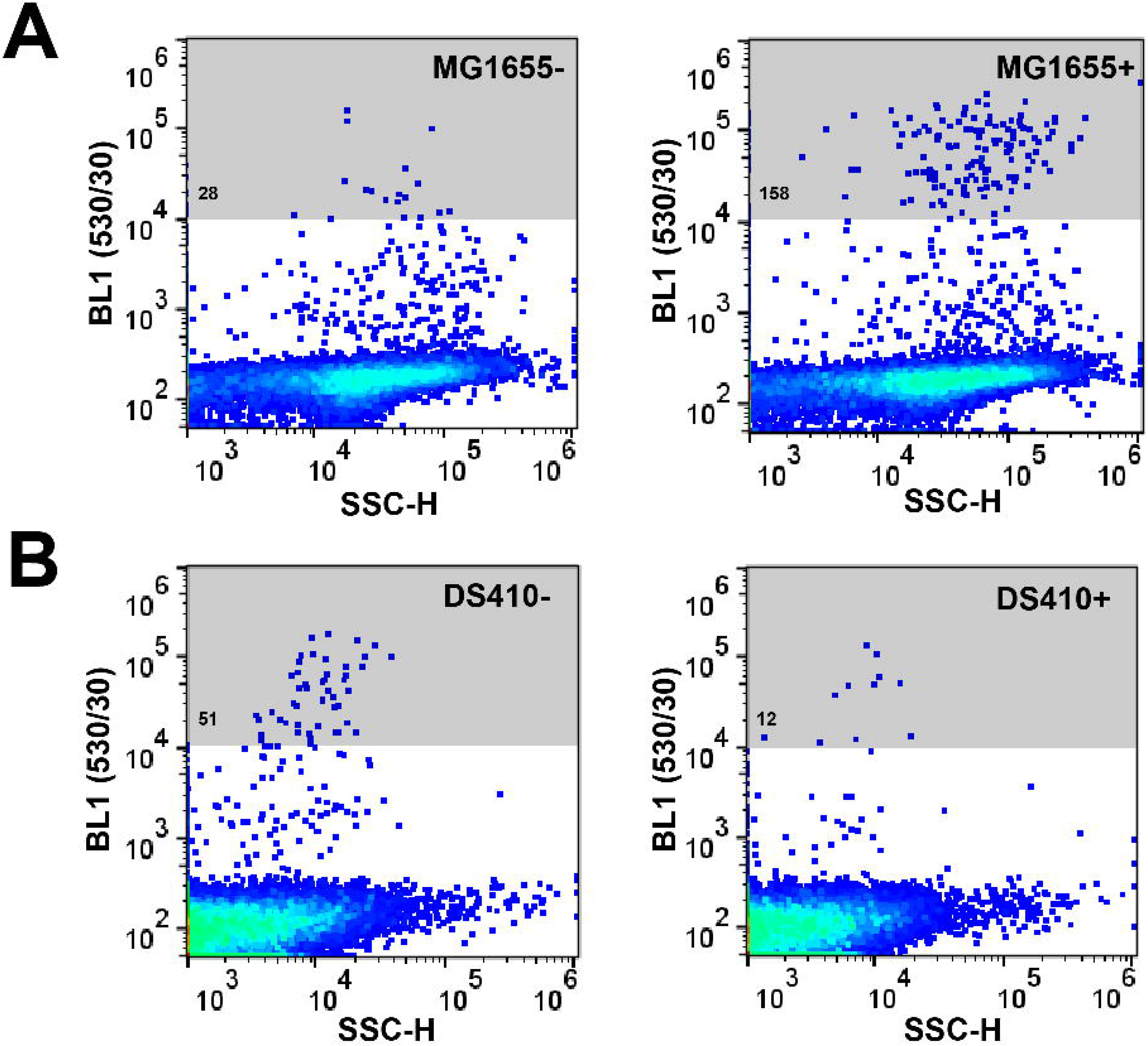
Standard curve for purified sfGFP fluorescence measurement on plate reader. Purified sfGFP fluorescence is measured on a 96-well plate reader at different dilutions at in 100 μL. Error bars are shown for three replicate measurements.

**Supplementary Figure 2.**
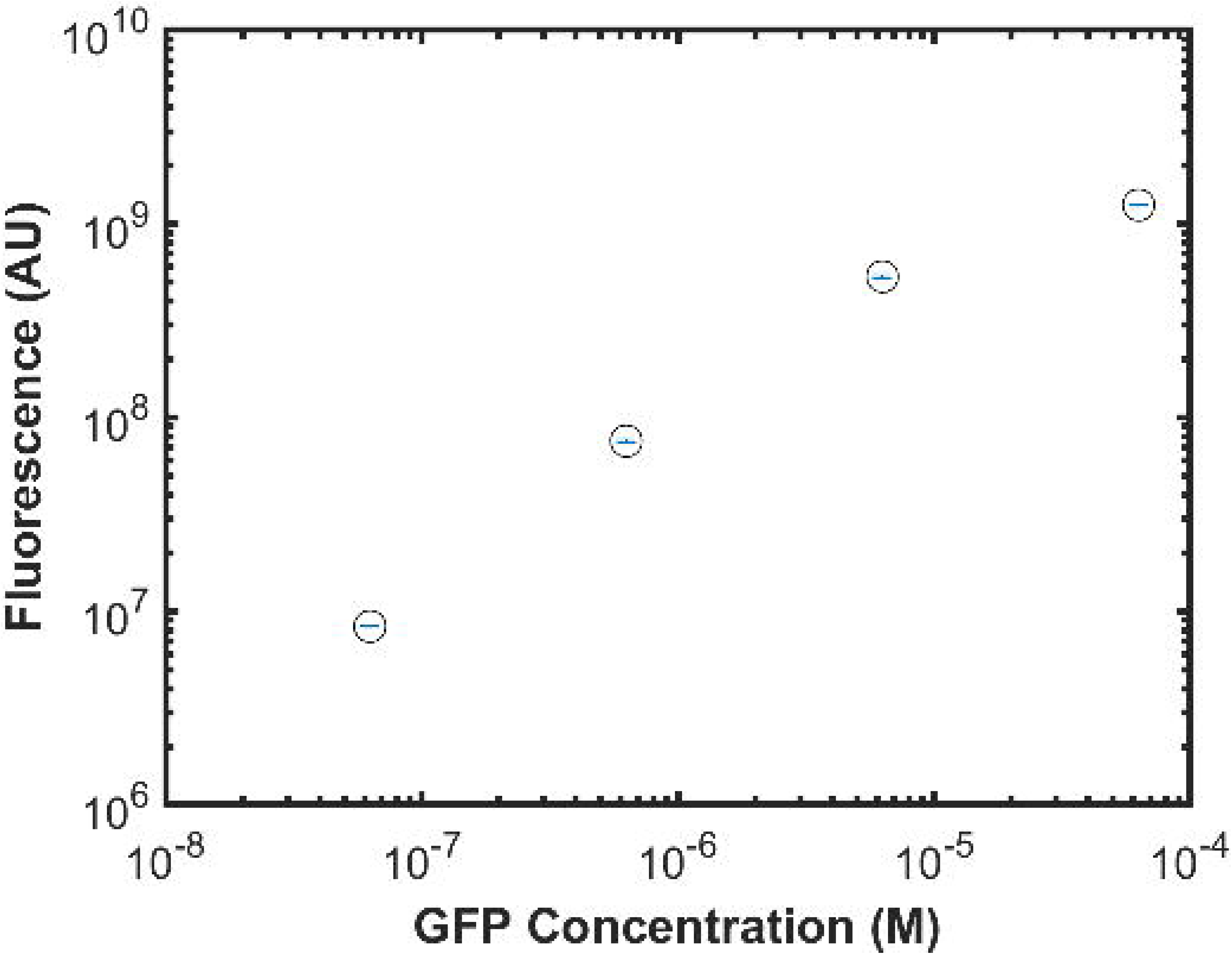
Plasmid transformation via electroporation of minicells could not be verified. Electroporation is performed on (A) wildtype *E. coli* and (B) purified DS410 minicells in the presence (+) and absence (−) of sfGFP plasmid. After a 2-hour incubation time for uptake and gene expression, minicells are sorted using flow cytometry measuring sfGFp fluorescence (530 nm emission with a bandwidth of 30 nm).

**Supplementary Video 1. Brightfield microscopy time lapse of minicell producing strain.**

Times given in hours:minutes.

**Supplementary Video 2. Fluorescence microscopy time lapse of minicell producing strain harboring sfGFP plasmid.** Times given in hours:minutes.

## Abbreviations

FACS: Fluorescence-activated cell sorting
sfGFP: Superfolder green fluorescent protein
LB: Luria-Bertani medium
OD: Optical density
PBS: Phosphate-buffered saline
SSC: Side scatter

## Acknowledgements

The authors would like to thank the CGSC for providing strain DS410, Atri Choksi and Pakpoom Subsoontorn for providing the sfGFP plasmid, Conary Meyer with contributions to establishing protocols and figures, and Endy Lab for discussions. This work was supported by the NSF Graduate Research Fellowship and Stanford Graduate Fellowship.

## Authors’ Contributions

DE and EW designed the experimental work and wrote the manuscript. EW performed microscopy of minicells and heterologous expression measurements. AJS performed FACS analysis of minicell purification and numerical minicell calculations. AJS and EW performed minicell purification, transformation, and electroporation experiment.

## References

1. Endy D. Foundations for engineering biology. Nature. 2005;438:449–453.

2. Benner SA, Sismour AM. Synthetic biology. Nature Reviews Genetics. 2005;6:533–543.

3. Keasling JD. Synthetic biology for synthetic chemistry. ACS Chemical Biology. 2008;3:64–76.

4. Khalil AS, Collins JJ. Synthetic biology: applications come of age. Nature Reviews Genetics. 2010;11:367–379.

5. Smanski MJ et al. Functional optimization of gene clusters by combinatorial design and assembly. Nature Biotechnology. 2014;32:1241–1249.

6. Bashor CJ, Collins JJ. Understanding biological regulation through synthetic biology. Annual Review of Biophysics. 2018;47:399–423.

7. Schmidt CM, Smolke CD. RNA switches for synthetic biology. Cold Spring Harbor Perspectives in Biology. 2019;11:1–11.

8. Carlson R. Estimating the biotech sector’s contribution to the US economy. Nature Biotechnology. 2016;34:247–55.

9. Aguilar A, Twardowski T, Wohlgemuth R. Bioeconomy for sustainable development Biotechnology Journal. 2019;14:e1800638.

10. Gibson DG, Benders GA, Andrews-Pfannkoch C, Denisova EA, Baden-Tillson H, Zaveri J, Stockwell TB, Brownley A, Thomas DW, Algire MA, et al. Complete chemical synthesis, assembly, and cloning of a *Mycoplasma genitalium* genome. Science. 2008:319:1215–1220.

11. Shao Y, Lu N, Wu Z, Cai C, Wang S, Zhang L-L, et al. Creating a functional single-chromosome yeast. Nature. 2018;560:331–335.

12. Carlson R. Competition and the future of reading and writing DNA. Synthetic Biology: Parts Devices and Applications. 2018;3–13.

13. Hutchison CA, Chuang R-Y, Noskov VN, Assad-Garcia N, Deerinck TJ, Ellisman MH, et al. Design and synthesis of a minimal bacterial genome. Science. 2016;351:aad6253.

14. Lluch-Senar M, Delgado J, Chen W-H, Lloréns-Rico V, O’Reilly FJ, Wodke JA, et al. Defining a minimal cell: essentiality of small ORFs and ncRNAs in a genome-reduced bacterium. Molecular Systems Biology. 2015;11:780.

15. Juhas M. On the road to synthetic life: the minimal cell and genome-scale engineering. Critical Reviews in Biotechnology. 2015;1–8.

16. Sung BH, Choe D, Kim SC, Cho B-K. Construction of a minimal genome as a chassis for synthetic biology. Essays In Biochemistry. 2016;60:337–346.

17. Baby V, Lachance J-C, Gagnon J, Lucier J-F, Matteau D, Knight T, Rodrigue S. Inferring the minimal genome of Mesoplasma florum by comparative genomics and transposon mutagenesis. 2018;3:1–14.

18. Breuer M, Earnest TM, Merryman C, Wise KS, Sun L, Michaela R, et al. Essential metabolism for a minimal Cell. eLife. 2019;8:e36842.

19. Glass JI, Merryman C, Wise KS, Hutchinson CA, Smith HO. Minimal cells — real and imagined. Cold Spring Harbor Perspectives in Biology. 2017;9:a023861.

20. Caschera F, Noireaux V. Compartmentalization of an all-*E. coli* cell-free expression system for the construction of a minimal cell. Artificial Life. 2016;2:185–195.

21. Stano P, Luisi PL. Semi-synthetic minimal cells: origin and recent developments. Current Opinion in Biotechnology. 2013;24:633–638.

22. Adamala KP, Martin-Alarcon DA, Guthrie-Honea KR, Boyden, ES. Engineering genetic circuit interactions within and between synthetic minimal cells. Nature Chemistry. 2017;9:431–439.

23. Jia H, Heymann M, Bernhard F, Schwille P, Kai L. Cell-free protein synthesis in micro compartments: building a minimal cell from biobricks. New Biotechnology. 2017;39:199–205.

24. Spoelstra WK, Deshpande S, Dekker C. Tailoring the appearance: what will synthetic cells look like? Current Opinion in Biotechnology. 2018;51:47–56.

25. Szostak JW, Bartel DP, Luisi PL. Synthesizing life. Nature. 2001;409:387–390.

26. Schwille P, Spatz J, Landfester K, Bodenschatz E, Herminghaus S, Sourjik V, et al. MaxSynBio: avenues towards creating cells from the bottom up. Angewandte Chemie. 2018;57:13382–13392.

27. Adler HI, Fisher WD, Cohen A, Hardigree AA. Miniature Escherichia coli cells deficient in DNA. Proceedings of the National Academy of Sciences of the United States of America. 1967;57:321–326.

28. Reeve JN, Mendelson NH, Coyne SI, Hallock LL, Cole RM. Minicells of *Bacillus subtilis*. Journal of Bacteriology. 1973;114:860–873.

29. Jaffé A, D’Ari R, Hiraga S. Minicell-forming mutants of *Escherichia coli*: production of minicells and anucleate rods. Journal of Bacteriology. 1988;170:3094–3101.

30. Ward Jr. JE, Lutkenhaus Joe. Overproduction of FtsZ induces minicell formation in *E. coli*. Cell. 1985;42:941–949.

31. Roozen KJ, Fenwick Jr RG, Curtiss III R. Synthesis of ribonucleic acid and protein in plasmid-containing minicells of Escherichia coli K-12. 1971;107:21–33.

32. Crooks H, Levy SB. Transcription of plasmid DNA in *Escherichia coli* minicells. Plasmid. 1983;72:66–72.

33. Dougan G, Kehoe M. The minicell system as a method for studying expression from plasmid DNA. Methods in Microbiology. 1984;17:233–258.

34. Dougan G, Fairweather NF. Detection of gene products expressed from plasmids. Methods in Microbiology. 1988;21:233–252.

35. Rampley CPN, Davison PA, Qian P, Preston GM, Hunter CN, Thompson IP, et al. Development of SimCells as a novel chassis for functional biosensors. Scientific Reports. 2017;7:1–10.

36. Giacalone MJ, Gentile AM, Lovitt BT, Xu T, Surber MW, Sabbadini RA. The use of bacterial minicells to transfer plasmid DNA to eukaryotic cells. Cellular Microbiology. 2006;8:1624–1633.

37. MacDiarmid JA, Mugridge NB, Weiss JC, Phillips L, Burn AL, Paulin RP, et al. Bacterially derived 400 nm particles for encapsulation and cancer cell targeting of chemotherapeutics. Cancer Cell. 2007;11:431–445.

38. MacDiarmid JA, Brahmbhatt H. Minicells: versatile vectors for targeted drug or si/shRNA cancer therapy. Current Opinion in Biotechnology. 2011;22:909–916.

39. Farley MM, Hu B, Margolin W, Liu J. Minicells, back in fashion. Journal of Bacteriology. 2016;198:1186–1195.

40. Levy, SB. Physical and functional characteristics of R-factor deoxyribonucleic acid segregated into *Escherichia coli* minicells. Journal of Bacteriology. 1971;108:300–308.

41. Shepherd N, Dennis PP, Bremer H. Cytoplasmic RNA polymerase in *Escherichia coli*. Journal of Bacteriology. 2001;183:2527–2534.

42. Levy, SB. Resistance of minicells to penicillin lysis: a method of obtaining large quantities of purified minicells. Journal of Bacteriology. 1970;103:836–839.

43. Jivrajani M, Shrivastava N, Nivsarkar M. A combination approach for rapid and high yielding purification of bacterial minicells. Journal of Microbiological Methods. 2013;92:340–343.

44. Bremer H, Dennis PP, Ehrenberg M. Free RNA polymerase and modeling global transcription in *Escherichia coli*. Biochimie. 2003;85:597–609.

45. Reeve JN. Bacteriophage infection of minicells. Molecular & General Genetics. 1977;79:73–79.

46. Ponta H, Reeve JN, Pfennig-Yeh M, Hirsch-Kauffman M, Schweiger M, Herrlich P. Productive T7 infection of *Escherichia* coli F^+^ cells and anucleate minicells. Nature. 1977;269:440–442.

47. García JA, Salas M. Bacteriophage φ29 infection of *Bacillus subtilis* minicells. Molecular & General Genetics. 1980;545:539–545.

48. Reeve JN, Lanka E, Schuster H. Synthesis of P1 ban protein in minicells infected by P1 mutants. Molecular & General Genetics. 1990;177:193–197.

49. Libby RT, Shaw JE, Reeve JN. Expression of coliphage T7 in aging anucleate minicells of *Escherichia Coli*. Mechanisms of Ageing and Development. 1984;27:197–206.

50. Bremer H, Dennis PP. Modulation of chemical composition and other parameters of the cell by growth rate. Escherichia coli and Salmonella: Cellular and Molecular Biology. ed. FC Neidhardt. 1996;1:1553–1569.

51. Kobayashi H. Regeneration of *Escherichia coli* from minicells through lateral gene transfer. Journal of Bacteriology. 2018;200:1–10.

52. Boulnois GJ, Timmis KN. Synthesis of plasmid-encoded polypeptides in maxicells. Advanced Molecular Genetics. 1984;204–207.

53. Gomaa AA, Klumpe HE, Luo ML, Selle K, Barrangou R, Beisel CL. Programmable removal of bacterial strains by use of genome-targeting CRISPR-Cas systems. mBio. 2014;5:e00928–13.

54. Caliando BJ, Voigt CA. Targeted DNA degradation using a CRISPR device stably carried in the host genome. Nature Communications. 2015;6:6989.

55. Schmitz-Esser S, Linka N, Collingro A, Beier CL, Neuhaus HE, Wagner M, Horn M. ATP/ADP translocases: a common feature of obligate intracellular amoebal symbionts related to *Chlamydiae* and *Rickettsiae*. Journal of Bacteriology. 2004;186:683–691.

56. Raymond ES, The Art of Unix Programming. Addison-Wesley. 2003.

57. AT&T Archives and History Center, UNIX: making computers easier to use. 2014. Retrieved from https://www.youtube.com/watch?v=XvDZLjaCJuw.

58. Antoine de Saint Exupéry. Citadelle. 1948.

59. Bonnet J, Subsoontorn P, Endy D. Rewritable digital data storage in live cells via engineered control of recombination directionality. Proceedings of the National Academy of Sciences of the United States of America. 2012;109:8884–8889.

60. Volkmer B, Heinemann M. Condition-dependent cell volume and concentration of *Escherichia coli* to facilitate data conversion for systems biology modeling. PloS ONE. 2011;6:e23126.

61. Edelstein AD et al. Advanced methods of microscope control using μManager software. Journal of Biological Methods. 2014;1:e11.

62. Balleza E, Kim JM, Cluzel P. Systematic characterization of maturation time of fluorescent proteins in living cells. Nature Methods. 2018;15:47–51.

